# Plant phylogeny and life history predict AM fungal species and genetic composition, but only life history and genetic composition predict feedback

**DOI:** 10.1101/2025.07.10.664051

**Authors:** Robert J. Ramos, Brianna L. Richards, Peggy A. Schultz, James D. Bever

## Abstract

Symbioses exert strong influences on host phenotypes; however, benefits from symbioses can increase or degrade over time. Understanding the context-dependence of reinforcing or degrading dynamics is pivotal to predicting stability of symbiotic benefits. Host phylogenetic relationships and host life history traits are two candidate axes that have been proposed to structure symbioses. However, the relative influence of host evolutionary history and life history on symbiont composition, and whether changes in symbiont composition translate into stronger mutualistic benefits is unknown. We tested the influence of plant phylogenetic relationships and plant life history on the composition of Arbuscular Mycorrhizal (AM) fungi, perhaps the most ancestral and influential of plant symbionts, and then tested whether AM fungal differentiation resulted in improved mutualism. We constructed mycobiomes composed of seven AM fungal isolates derived from tallgrass prairie and grew them for two growing seasons with 38 grassland plant species. We found that host phylogenetic structure was a significant predictor of the composition of AM fungal communities and the genetic composition of AM fungal species, patterns consistent with phylosymbiosis. However, the phylogenetic structure of AM fungi failed to translate to improved benefit to their host. While AM fungi generally improved plant growth and mycorrhizal feedback was generally positive, the strength of feedback was not predicted by plant phylogenetic distance. The composition of the AM fungal community and genetic composition within AM fungal species were also significantly influenced by plant life history. In addition, feedbacks between early and late successional species were generally positive. Interestingly, changes in species composition did not predict feedback, while changes in genetic composition of the two most abundant AM fungal species predicted positive mycorrhizal feedback. These reinforcing mycorrhizal feedbacks across life history can mediate plant species turnover during succession and suggests that consideration of mycorrhizal dynamics could improve restoration.

## Introduction

Symbioses exert strong influences on phenotypes. Symbionts are particularly important to plants, with 80% of all terrestrial plant species, including most crop plants, associating with arbuscular mycorrhizal (AM) fungi [1]. This symbiosis has been identified as ancestral to all root symbioses [2] with arbuscules predating the origin of roots [3,4]. AM fungi mediate host acquisition of soil resources, particularly phosphorus, and can have a large influence on plant community dynamics [5–9]. Together, these aspects of AM fungi illustrate the critical importance of symbionts on host evolution and ecology, the understanding of which is a central question in biology.

Host phylogeny has been posited as an important factor structuring symbiont composition and function [10]. Phylogenetic structure of symbiont composition, either of symbiont species or genetic variants within symbiont species, has been called phylosymbiosis and has been observed in many systems including: the internal microbiomes of insects (*Drosophila* flies, mosquitoes, and *Nasonia* wasps) [11–13], the gut microbiota of animals (*Peromyscus* deer mice, pack rats, house mice, cranes, decapod shrimps) [11,14–16], and the bacterial communities of corals [17,18], providing a basis for generalizing across host-symbiont interactions. To date, there has been no comprehensive test of plant phylogenetic influence on AM fungal fitness and composition.

Inference on symbiont function from observations of phylogenetic structure in symbiont composition are challenging as they depend upon differential influence of symbionts on host fitness. While it is tempting to assume that differentiation in symbiont composition results in improved symbiont functioning for their hosts, fitness effects from host-specific differentiation of symbionts can result in positive or negative feedback [19] which correspond to strengthening or deteriorating host fitness with phylosymbiosis. Several studies have found evidence of reinforcing coadaptation of symbionts on their hosts, including within *Nasonia* wasp species [12,13] and within some *Peromyscus* mouse species, but not others [11]. The benefit of AM fungi to their hosts has been shown to decline [20,21] or increase [22,23] with host-specific differentiation of symbionts. Whether host phylogenetics influence the likelihood of positive, reinforcing feedback remains an open question.

It is also possible that host traits unrelated to phylogeny are primary factors structuring symbiont differentiation on hosts. While the candidate host traits driving symbiont differentiation are many, major axis of plant trait variation has been shown to align with the fast-slow/acquisitive-conservative continuum of life history strategies [24,25]. Plant life history has also been shown to be important for plant-AM fungal interactions. Specifically, slow growing late successional plant species have been shown to be more responsive to AM fungi [26,27], to be more sensitive to AM fungal identity [28], and to benefit particularly from AM fungi found in undisturbed sites dominated by late successional plant species [29,30], potentially generating positive feedback [23]. Such trait mediated AM fungal feedbacks could have important consequences for community coexistence and turnover during succession.

Plants acquire their AM fungal symbionts from the environment each generation—there is no horizontal inheritance of this symbiosis. While AM fungal species have been shown to be functionally variable [8,31], variants within AM fungal species can also vary in their ecology [32,33]. AM fungi are multinucleated, and different nuclei within AM fungal individuals can have substantially different genetic content [34–36]. Parent AM fungal cells can segregate nuclei, passing different nuclei to different offspring, and these segregants have been shown to differ in their impacts on host plants [32,33,37–39]. To date, we do not know the relative importance of changes in species composition vs genetic composition within species in driving functional consequences of AM fungal dynamics.

To test how plant host identity, plant host phylogenetic relationships, and plant life history characteristics shape symbiotic AM fungal communities, we grew a common community of AM fungi, a constructed synthetic mycobiome, with 38 individual plant species (Fig. 1a) representing a phylogenetically diverse group of early and late successional host prairie plants (S1 Fig and S2 Table). All plants were grown in a common greenhouse environment to remove potential confounding environmental effects [10]. We monitored the changes in AM fungal composition over two growing seasons using LSU rDNA amplicon sequencing [40–42]. As rDNA of AM fungi are highly polymorphic [43,44], we grouped amplicon sequence variants (ASVs) into species to permit assessment of changes in AM fungal species composition and changes in genetic composition of individual AM fungal species (Fig. 1b). We tested for consistent effects across plant phylogeny and across plant life history categories with two approaches. PerMANOVA controls for non-independence of amplicons in individual samples. Phylogenetic mixed models explicitly test for host phylogeny influence on AM fungal composition and simultaneously control for phylogenetic non-independence. Both test for effects of host life history on AM fungal composition (Fig. 1c).

**Fig 1:**
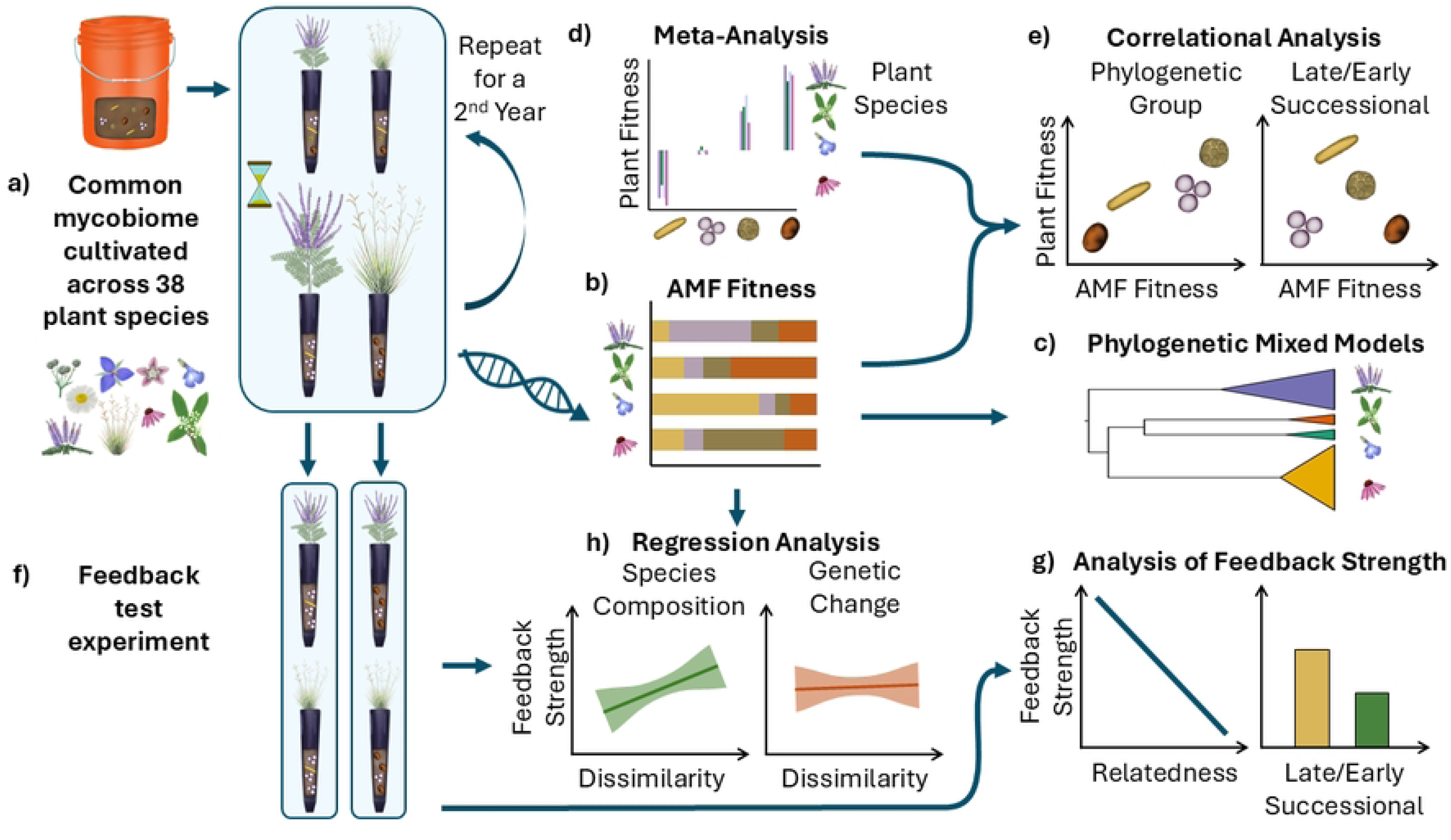
Conceptual Figure of the Experimental Design. A common AM fungal community composed of mixtures of characterized isolates was grown with 38 plant species for two growing seasons (a). Changes in relative abundance of AM fungi were monitored using amplicon sequencing (b) and used to test for plant phylogenetic and life history impacts on AM fungal species and genetic composition (c). The effect of the AM fungal inoculum strains used in this experiment on plant species was characterized using meta-analyses of prior studies (d). Correlations of AM fungal relative fitness and fungal impacts (e) on host phylogenetic and life history groupings were tested as one measure: mycorrhizal feedback [19]. Feedback was also measured directly by testing growth responses of different plant species to trained AM fungal communities (f). Pairwise feedbacks, as represented by interaction coefficients (Bever et al. 1997), were calculated and used to test patterns of strength of feedback across host phylogeny and life history (g). We also tested whether the observed changes in AM fungal species composition or changes in genetic composition within species predicted strength of mycorrhizal feedback (h).

We used multiple approaches to assess the ecological impact of changes in AM fungal composition (Fig 1). As our constructed mycobiomes included isolates that have been characterized in their impacts on host growth across plant family and across plant life history [28,45], we test for correlations of AM fungal fitness responses to hosts with AM fungal impacts on host growth (Fig. 1d,e,). A positive correlation of host and symbiont fitness impacts predicts positive, reinforcing feedback that would generate co-adaptation [19,46]. Secondly, we directly tested the functional consequences of host-specific differentiation in AM mycobiome composition using a feedback assay (Fig. 1f) in which plant species were grown with the differentiated mycobiomes in full factorial combinations [47,48]. We then tested whether the strength of feedback is positively correlated with phylogenetic distance, as expected by co-adapted phylosymbiosis, and whether AM fungal feedback varies consistently with plant life history, as expected from mycorrhizal dynamics influencing species turnover during succession (Fig. 1g). Finally, we tested whether changes in AM fungal species or genetic composition within AM fungal species predicted resulting feedbacks (Fig. 1h).

## Results

We found significant influence of plant phylogeny and plant life history on the species and genetic composition of the AM mycobiome. This holds true whether controlling for the multivariate nature of the amplicon data using PerMANOVA or whether testing for and controlling for phylogenetic relationships of host plants using phylogenetic generalized mixed models (PGLMMs). As these results are complementary, but redundant, we present the PGLMM results which provide a direct test of phylosymbiosis below and the PerMANOVA within the appendix (S3 and S4 Tables).

### Host phylogeny impacts on AM Mycobiome

Grouping ASV counts by individual AM fungal species, we found significant changes in AM fungal composition and density with plant phylogeny. Host phylogeny significantly affected AM fungal diversity and density in year 1 (p<0.01, p<0.001) and the relative proportion of *Ce. pellucida* in year 1 (p<0.05), *R. fulgida* in year 2 (p<0.001), and *E. infrequens* in both years (p<0.001, p<0.05, S5 Table, Fig 2). This suggests that phylogenetically proximate plant species have more similar AM fungal composition, reflecting more similar impacts of host on AM fungal relative fitness, than more phylogenetically distant plant species. *Ce. pellucida* had high relative growth rates with plants in Apocynaceae, *E. infrequens* had low relative fitness with grasses, and *R. fulgida* tended to perform best with grass species (Fig 2).

**Fig 2:**
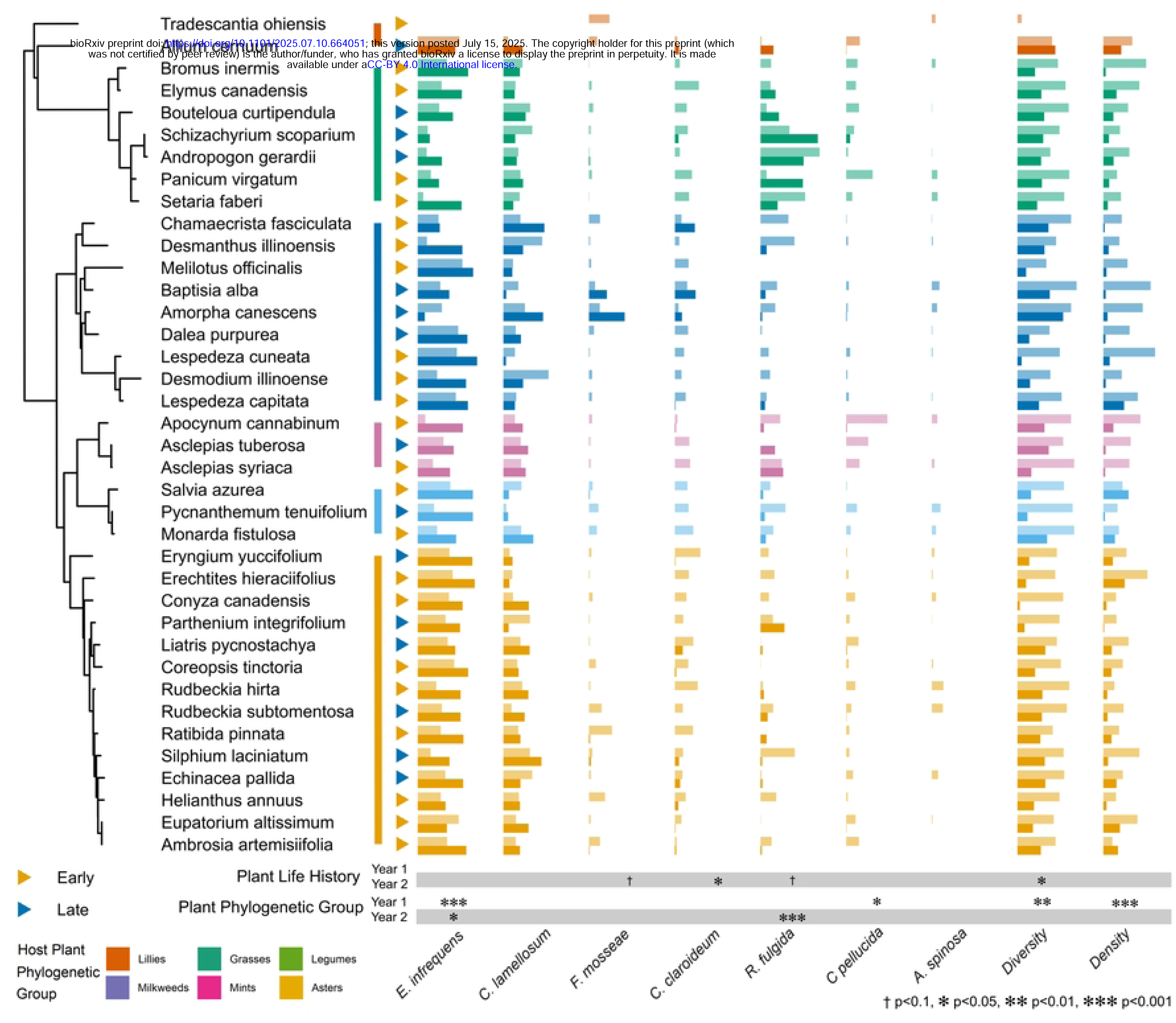
AM Fungal Species Relative Abundances Across Plant Phylogeny. AM fungal relative abundances for *Cl. lamellosum*, *Cl. claroideum*, *E. infrequens*, *F. mosseae*, *R. fulgida*, *Ce. pellucida*, and *A. spinosa* are presented alongside estimated abundance (proportion of ASVs identified as inoculated over total AM fungal ASVs) and Shannon’s Diversity. Results are by host plant species arranged phylogenetically. Significance phylogenetic effects are calculated with a Likelihood Ratio Test using the Chi square distribution. Year 1 and Year 2 data are presented as grouped columns with year one being the lighter shaded top bar.

### Host phylogeny impacts on genetic composition of AM fungal species

We also tested the influence of host phylogeny on the genetic composition of individual AM fungal species. ASV’s identified for each species were tested in order of decreasing overall abundance. We found significant differences in ASV composition due to plant phylogenetic relatedness for the most abundant *E. infrequens* ASVs for both years (p<0.01, Fig 3). The most abundant ASV of *Cl. lamellosum* was marginally affected by plant phylogeny in year 1 (p>0.1). *F. mosseae’s*, *Cl. claroideum’s,* and *Ce. pellucida’s* third most abundant ASV had a significant effect of host plant phylogenetic distance in year 1 (p<0.01, p<0.001, p<0.001). Finally, *R. fulgida* was marginally affected by phylogeny for its seventh most abundant ASV in year 1 (p>0.1). There were also significant effects of phylogeny for its most abundant ASV in year 2 (p>0.001).

**Fig 3:**
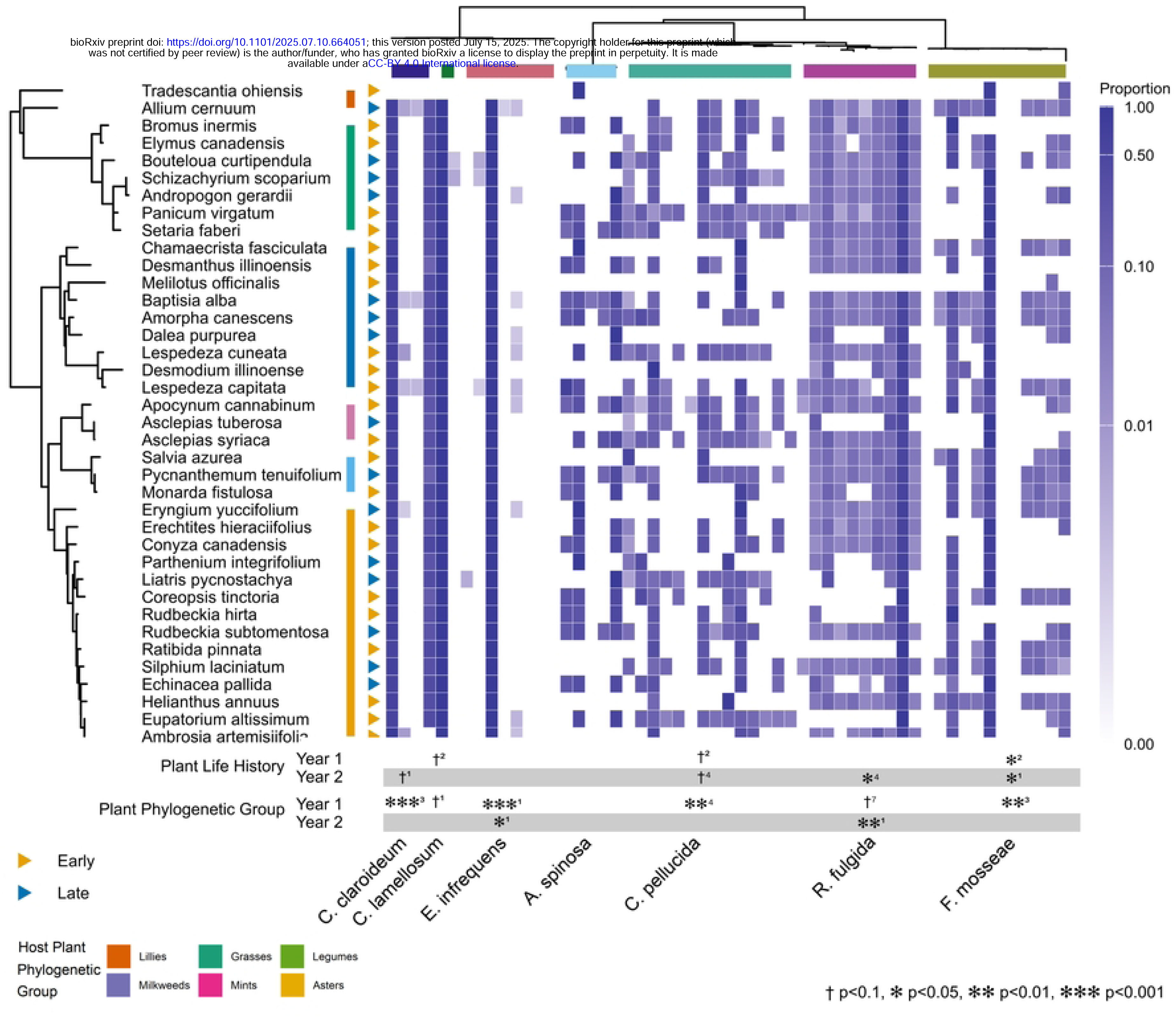
AM Fungal Genetic Variation Across Plant Phylogeny. AM fungal genetic variation (ASV relative abundances) were tested for significant effects of phylogenetic distance within *Cl. lamellosum*, *Cl. claroideum*, *E. infrequens*, *F. mosseae*, *R. fulgida*, *Ce. pellucida*, and *A. spinosa*. The significance of each phylogenetic effect is calculated with a Likelihood Ratio Test using the Chi square distribution. Results are reported for the most common ASV with a significant result. The rank of the most common significant ASV is given as a superscript next to the significance symbols.

### Plant life history impacts on AM mycobiome

PGLMMs of total ASV counts grouped by individual AM fungal species showed significant changes in AM fungal composition with plant life history. The relative abundance of *F. mosseae* and *R. fulgida* were marginally significantly higher when associated with late successional plants in year 2, with *Cl. claroideum* being significantly higher (p<0.1, p<0.1, p<0.05, S5 Table, Fig 2). Late successional plant species had more diverse AM fungal communities than early successional plant species (Shannon index, year 2 p≤0.05, S6 Fig). Estimated AM fungal density was not significantly affected by host plant life history in either year (p>0.1).

### Plant life history impacts on genetic composition of AM fungal species

Host plant life history significantly impacted the genetic composition of several AM fungal species. PGLMM analysis found the relative abundance of the second most abundant ASV of *Cl. lamellosum*, *F. mosseae*, and *Ce. pellucida* were each affected by host plant life history in year 1 (p<0.1, p>0.05, p>0.1, Fig 3). In year 2, the most abundant *F. mosseae* ASV and fourth most abundant *Ce. pellucida* ASV were significantly affected by host plant life history (p<0.05, p>0.05). Finally, the most abundant *Cl. claroideum* ASV had a marginal response to host plant life in year 2 only, and the fourth most abundant *F. mosseae* ASV had a significant response, (p<0.1, p<0.05).

### Correlation of host and symbiont relative fitness

Previous work with the AM fungal isolates used in this experiment [28,45] allowed us to test for general patterns across four family groups: Asters (Asteraceae), Legumes (Fabaceae), Lillies (Amaryllidaceae and Commelinaceae), and Grasses (Poaceae). We were particularly interested in the differential impacts on plant families and their correlation with the differential fitness responses to these families. We found that *E. infrequens*, *Cl. lamellosum*, and *Cl. claroideum* confer greater benefits to plants in the Asters relative to plants in the Grasses group (S7B Fig). We also saw that *E. infrequens*, *Cl. lamellosum*, *Cl. claroideum, F. mosseae, A. spinosa,* and *Ce. pellucida* confer greater benefits to plants in the Lilly group compared to the Grasses. These effects are weakly positively correlated with the relative fitness responses to these plant families (Fig 4). This provided weak evidence of mycorrhizal positive feedback particularly between plants in the comparison of the phylogenetic groups Asters and Grasses (R^2^_yr1_=0.32, p=0.187, R^2^_yr2_=0.48, p=0.127) as compared to Legumes – Grasses (R^2^ =0.32, p=0.188, R^2^ =0.49, p=0.183) and Lillies – Grasses (R^2^ =0.03, p=0.702, R^2^ =0.01, p=0.877). This may suggest a strengthening of host fitness with phylosymbiosis, though none of these correlations were significant.

**Fig 4:**
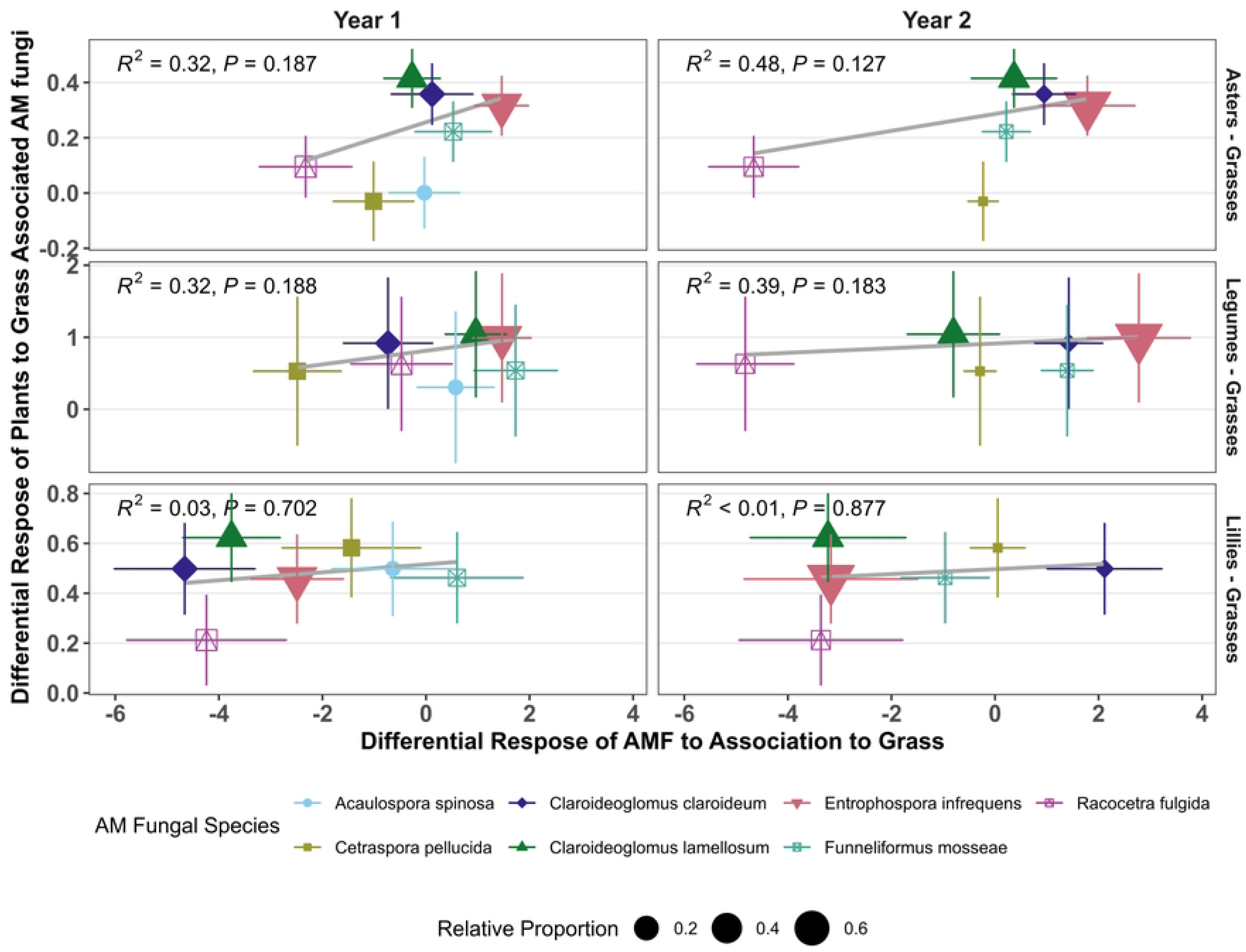
Predicted Feedbacks Between Phylogenetic Groups. We analyzed the correlation of meta-analysis results of host plant response to AM fungal species associated with host plants of different phylogenetic groups, with the measured differential AM fungal response to host plant phylogenetic group. R^2^ and p values corresponded to linear correlations weighted by AM fungal relative abundance. There were no significant correlations at the p≤0.1 level.

We also found the growth promotion by AM fungal isolates differed in early vs. late successional plant species (S7A Fig). Moreover, there was a negative correlation between AM fungal growth and benefits to plant hosts, with late successional plants growing less well with the AM fungal isolates grown on other late successional hosts (R^2^ =0.23, p=0.274, R^2^ =0.94, p=0.001, Fig 5). This pattern is consistent with changes in AM fungal composition generating negative feedback between plants of different life history strategies. Given that AM fungal species diversity increased with late successional plant species in the second year of AM mycobiome training (Fig 2), it is possible that the change in AM fungal diversity feedback on fitness of late successional plant species. However, when analyzing two studies comparing early and late successional plant species in response to single and mixed inocula, we did not find a significant difference in response to AM fungal diversity between plant life history stages (p= 0.748, S8 Fig), casting doubt on feedback through changes in mycorrhizal diversity.

**Fig 5:**
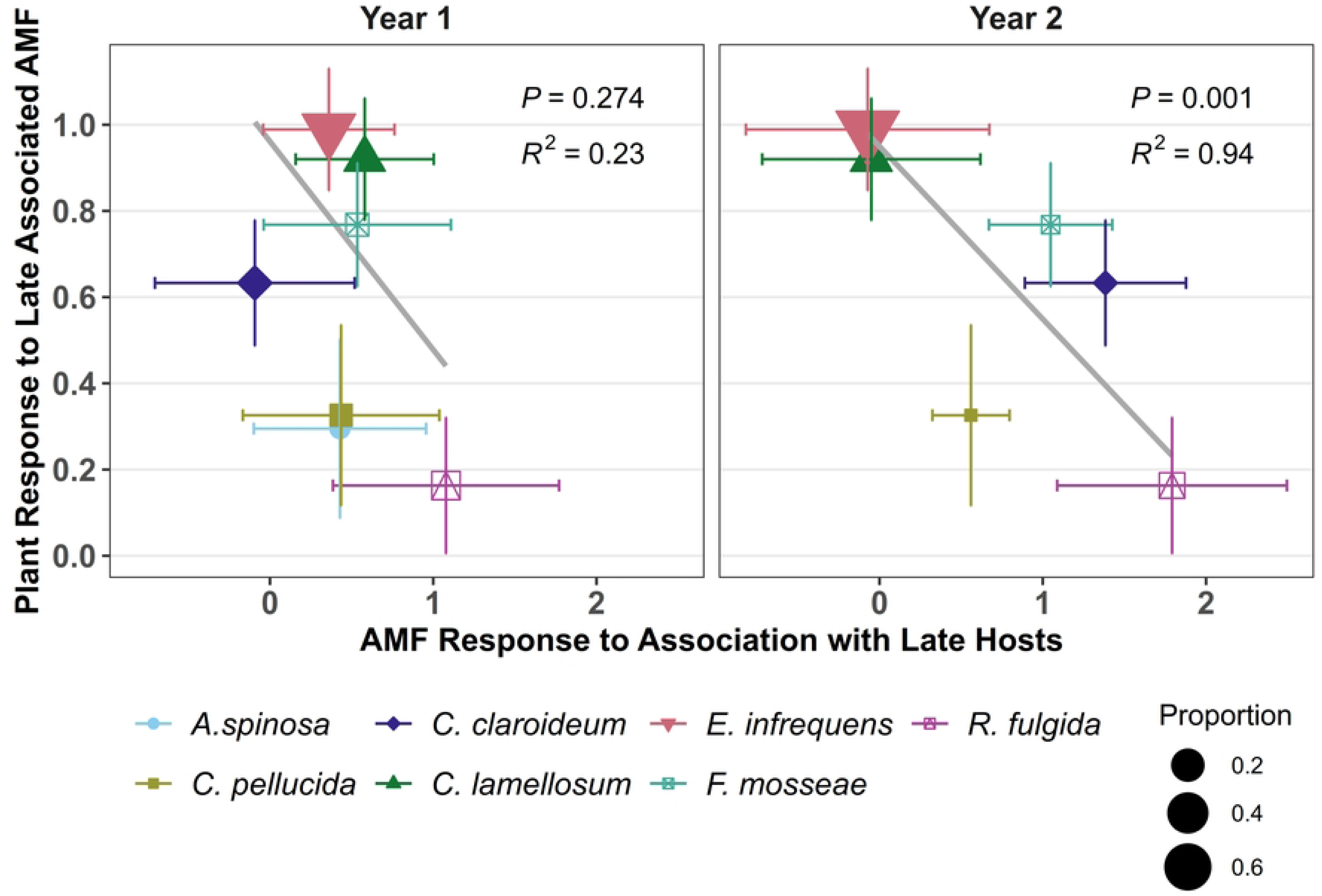
Predicted Feedbacks Between Life History Categories. We analyzed the correlation of meta-analysis results of host plant response to AM fungal species associated with host plants of different life history categories to the measured differential AM fungal response to host plant life history category. R^2^ and p values correspond to linear correlations weighted by AM fungal relative abundance. There is a significant positive correlation in year 2 (p=0.001).

### AM fungal feedbacks: tests of patterns

AM fungal feedback between pairs of plant species varied from significantly positive to significantly negative (Fig 6). Across all pairs of plants, median (p = 0.003) and mean (t = 3.9472, p≤ 0.001) AM mycobiome feedback was generally positive (S9 Fig). However, there was no relationship between strength of feedback and plant phylogenetic distance. The best fit was a marginal quadratic fit (x: p=0.167, x^2^: p=0.085, Fig 6A), as closely related species and most distantly related species tended to have zero or negative feedbacks. Together this indicates that the observed plant phylogenetic influence on AM mycobiome composition did not consistently result in improved host fitness.

**Fig 6:**
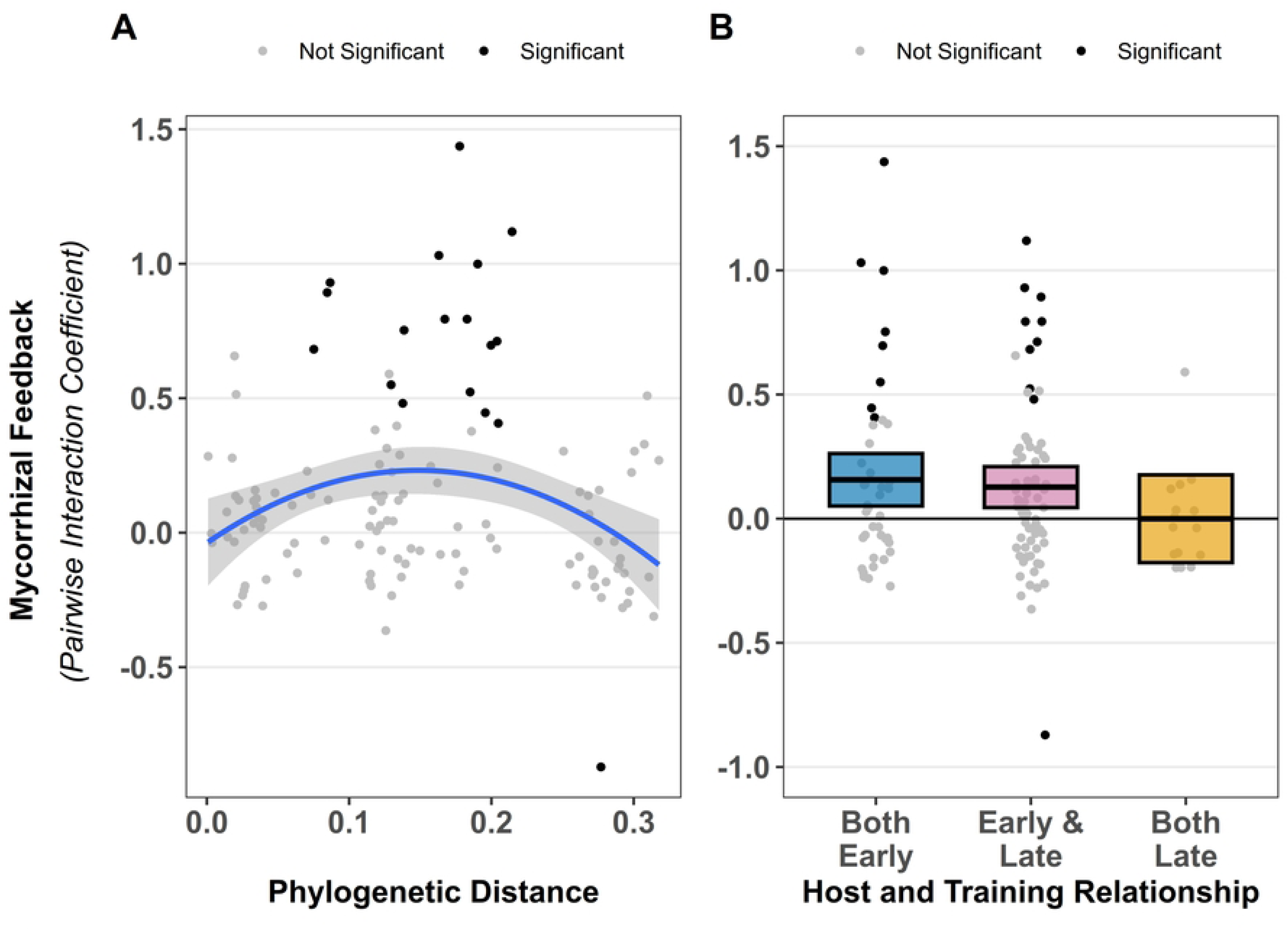
Measured Feedback Strength Between AM fungi and Plant Hosts. Mycorrhizal feedback, measured as pairwise interaction coefficients, are plotted on the Y-axis in all plots. Phylogenetic distance (A) had only a marginally significant quadratic relationship with fitness (p=0.085). (B) shows feedbacks broken out by both early, both late, and early and late host plant parings. Both early (p=0.011) and early-late (0.003) parings had significantly positive feedbacks. Both panels show individual pairwise interaction values plotted as shaded dots, with grey for insignificant interaction coefficients and black for significant interactions (p>0.05).

When analyzing feedback effects between plant families (restricted to the three best represented families: grasses, asters, and legumes). We tended to find mean positive feedback between legume species (t_13_=2.233, p=0.058), negative feedback between Aster–Grass pairings (t_18_=-2.236, p=0.029) and Grass–Grass pairings (t_6_=-2.082, p=0.082). Other comparisons were not consistently different from zero (S10 Fig). Median Legume-Legume pairings remained marginally positive (p-value = 0.078), and Aster-Grass remained significantly negative (p=0.014). Grass–Grass pairings did not have significant negative median feedbacks (p=0.109).

In comparing within and between plant life history categories, the median pairwise feedbacks between pairs of early successional species tended to be positive (p=0.069), and the mean feedback was significantly positive (t_41_=2.654, p=0.01, Fig 6B). Comparing feedback between early and late successional plant species, both the median (p=0.009) and mean values (t_68_=3.055, p=0.003) were significantly positive. When both plant species were late successional neither the median (p=0.488) nor the mean (t_14_=-0.009, p=0.993) feedback were significantly different from zero.

### AM fungal feedbacks: community and/or genetic drivers

Differences in AM fungal species composition and/or differences in genetic composition within AM fungal species could drive mycorrhizal feedbacks. Changes in AM fungal species composition, as represented by dissimilarity between plant species pairs, failed to predict feedback strength (Fig 7a, R^2^=0, F= 0.2922, p=0.59). However, changes in genetic composition of individual AM fungal species did predict mycorrhizal feedback. Specifically, positive pairwise feedback was significantly predicted by dissimilarity in genetic composition of the two most common AM fungal species: *E. infrequens* (Fig 7b, R^2^=0.13, F= 17.78, p<0.001) and *Cl. lamellosum* (Fig 7c, R^2^=0.1, F=13.27, p<0.001). Genetic dissimilarity of other AM fungal species did not predict pairwise feedback (S11 Fig).

**Fig 7:**
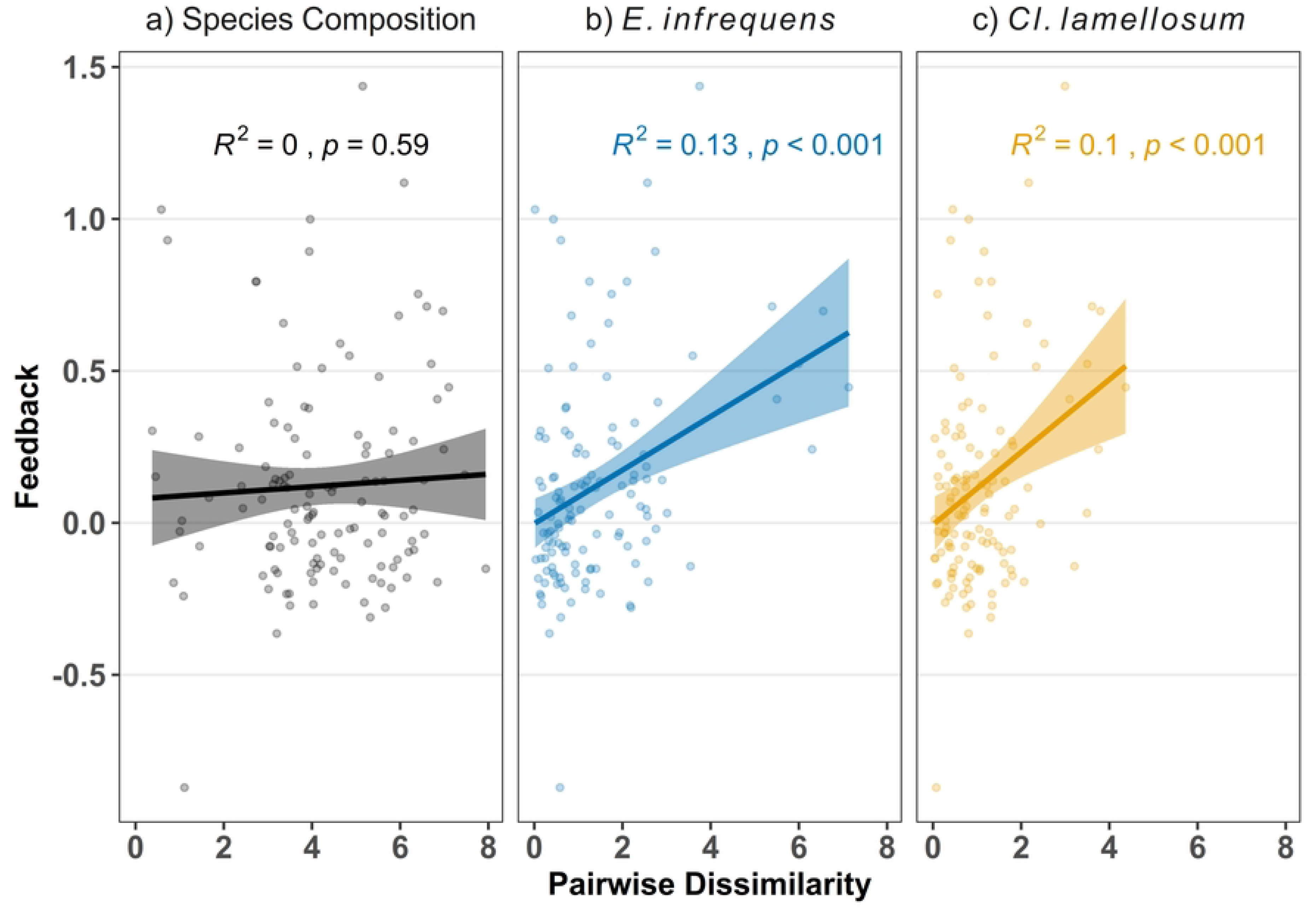
Mycorrhizal Feedback Strength Predicted by Genetic Dissimilarity of AM fungi. Regressions of strength of pairwise feedback against measures of AM fungal dissimilarity. AM fungal species composition, as measured by pairwise dissimilarity of total species counts (Aitchison distance) did not predict feedbacks (p=0.59). Pairwise feedback was significantly predicted by genetic dissimilarity of two AM fungal species, *E. infrequens* (p<0.001) and *Cl. lamellosum* (p<0.001).

## Discussion

Using 38 plant species, we present a uniquely robust test of plant phylogeny and life history impacts on the composition and symbiotic function of arbuscular mycorrhizal (AM) fungi. We found strong phylosymbiosis, as phylogenetic distance was an important predictor of the differences in AM fungal diversity, density, and community composition across plant host species. In addition, we detected significant differences in genetic composition of individual AM fungal species across plant host species. These differences are often predicted by plant phylogenetic distance, indicating that AM fungal species can evolve rapidly in response to their host. However, while AM fungi generally improved host growth, we found only weak evidence that the phylogenetically structured divergence of symbiont community and genetic composition consistently altered symbiont impacts on their hosts. After controlling for the effect of plant phylogeny, we also found that AM fungal density, community diversity, community composition, and genetic composition within species changed with plant life history, as represented by early vs late successional stage categories. In this case, the shift in symbiont composition with plant life history differentially benefited plants of the same life history generating positive mycorrhizal feedback. Differential positive feedbacks that vary between plants of different with life history characteristics could contribute to plant species turn over during succession. Interestingly, our results suggest that the genetic change within AM fungal species, rather than changes in relative abundance of AM fungal species, generated the observed positive mycorrhizal feedback. Together, our results indicate that both plant phylogeny and plant life history traits govern the associated AM fungal community and genetic composition, but only plant life history traits have strong evidence for consistent reinforcing positive feedbacks on plant fitness.

We were able to confirm that plant host identity and phylogenetic relationships were important in shaping the AM fungal community composition. Previous studies have demonstrated that AM fungal growth rates varied between individual plant species [48,49]. We affirm that this result is general across plant species, as AM fungal composition diverged significantly across the 38 plant species studied. Moreover, host-specific differentiation of AM fungal composition had a strong phylogenetic signature. That is, the growth rates of individual AM fungal species tended to be similar on phylogenetically similar host plants. This phylogenetic influence was evident in the differential abundance of *E. infrequens, R. fulgida*, and *Ce. pellucida* (Fig 2). Phylogenetic distance also predicted measures of AM fungal diversity. While previous work has shown phylogenetic signature to plant species specific differentiation of soil bacterial and fungal communities [50], this study provides clear evidence of phylogenetic structure to host-specific differentiation of AM fungal communities.

We note that the influence of host identity on AM fungal composition does not appear to result from differential exclusion of AM fungal species from roots, as would be expected from incompatibilities of plant-AM fungal signaling. Rather, inoculated AM fungi were found with all plant species. Given that inoculated AM fungi had equivalent densities across each plant species at the beginning of the experiment, the change in AM fungal composition is due to host-specific differences in AM fungal growth rates (i.e., fitness). The observation of substantial influence of host phylogeny on AM fungal composition may reflect that plant influence on AM fungal relative fitness can be moderated by phylogenetically conserved plant traits, such as root architecture [51] or secondary chemical composition [52].

Our first measure of host phylogenetic influence on mycobiome composition solely examined species level changes, and this may underestimate the impact of host phylogeny which could operate on genetic changes within each species. AM fungal species and isolates have highly variable rDNA genes [43,53], and we observed changes in rDNA composition with plant host species across most AM fungal species included in this study. Using both PerMANOVA and PGLMM methods, we detected unique host family/phylogenetic structure of the genetic composition of all our AM fungal species with the exception of *A. spinosa*, a species that had very low relative abundances. Our most abundant AM fungal species, *E. infequens,* exhibited host phylogenetic influence on its genetic composition for both years, while *Cl. lammelosum*, *Cl. claroideum, Ce. pellucida*, *R. fulgida*, and *F. mosseae* exhibited phylogenetic responses in genetic composition in one year regardless of the analysis used. In addition, host plant life history showed at least a marginal effect on AM fungal genetic composition for all species except *E. infrequens* and *A. spinosa*. Previous work had shown that genetic composition of AM fungal species can change rapidly with environment and host plant species [33,54,55]. These rapid changes could be due to differential segregation of nuclei in multi-nucleate cells [32,33,38]. We provide evidence that such genetic change can occur widely across AM fungal taxa and that the host-specific effects on AM fungal genetic composition are structured by plant phylogeny.

We find host-specific changes in AM fungi composition generate feedback on plant growth that is, on average, positive and ranges from significantly negative to strongly positive for individual plant species pairs. This is consistent with previous studies [20,22,23]. Our study provides the first attempt to generalize the patterns of AM fungal feedback, specifically testing whether AM fungal feedback is structured by plant phylogeny or plant life history. We test these patterns by testing correlations of AM fungal differentiation with growth impacts of these fungi observed in previous growth assays with our AM fungal isolates [28,45]. In addition to this correlational approach, we also measured AM fungal feedback effects by analyzing patterns observed among feedbacks directly measured using an independent greenhouse assay [48]. While the correlation of plant and AM fungal fitness responses only considers the impacts of changes in AM fungal species composition, the direct measurements of AM fungal feedbacks integrates across changes in AM fungal species and genetic composition.

Both correlations of relative plant and fungal fitness (Fig 4) and patterns of AM fungal feedbacks (Fig 6C) show weak responses to plant phylogenetic structure. That is, the AM fungi with the highest growth rates on a particular group of phylogenetically-related plants do not generally promote the growth of these phylogenetically-related plants better than other AM fungi. Interestingly, the best fit of AM fungal feedbacks to plant phylogenetic relatedness was curvilinear, with regions of phylogenetic distance over which plants tended to benefit more from AM fungal communities cultivated by more distantly related host plants and regions where plants tended to benefit more from AM fungal communities cultivated by more phylogenetically similar host plants (Fig 6C). Given that this is the first test of such relationships, more work is necessary in the plant-AM fungal systems as well as other host-symbiont interactions.

In contrast to plant phylogeny, AM fungal feedbacks are structured by plant life history. AM fungal feedback, as measured by direct greenhouse assays, were consistently positive between early and late successional plant species (Fig 6b). This result is consistent with observations greater responsiveness of late successional species to mycorrhizal fungi [28,45], and of positive mycorrhizal feedback observed in mesocosms [23] and field inoculations with prairie AM fungi [31,56,57]. Positive mycorrhizal feedback could play an important role in plant dynamics during succession and restoration, potentially accelerating turnover during succession, or contribute to alternative stable states [19]. Mycorrhizal feedback between pairs of late successional plant species were not generally different from zero, consistent with all late successional species benefiting equivalently from changes in mycorrhizal composition, while feedback between pairs of early successional species tended to be positive, suggesting that this life-history category includes plants with substantial variation in relationships with AM fungi.

The change in AM fungal species composition was not sufficient to explain the positive AM fungal feedback between early and late successional plant species (Fig. 5). In fact, across all the data, the change in AM fungal species composition did not predict observed patterns of mycorrhizal feedback (Fig. 7a). AM fungal species diversity also increased with late successional plant species, and AM fungal diversity can influence host response [58] potentially generating positive AM fungal feedbacks. However, this hypothesis was not supported by meta-analysis of differential benefits to late successional plant species from AM fungal diversity (S8 Fig).

The observed patterns of mycorrhizal feedback appear to have resulted from changes in genetic composition within individual AM fungal species. This hypothesis is directly supported as the genetic change within the two most common and beneficial AM fungal species, *E. infrequens* and *Cl. Lamellosum*, significantly predicted the strength of positive mycorrhizal feedback (Fig. 7b,c). Our results are consistent with previous observation of host-driven genetic changes within AM fungal isolates altering AM fungal impacts on plant fitness [32,38], and of patterns consistent with a role of genetic variation within AM fungal species generating mycorrhizal feedback [21]. Our work illustrates that rapid genetic change within AM fungal species can have functional consequences, even exceeding those of changes in AM fungal species composition. Our result highlights limitations to efforts to understand AM fungal ecology using patterns of species level morphological traits [59,60] as we find that mycorrhizal feedback, the product of variation in growth responses to hosts and impacts on hosts, meaningfully varied between genotypes within AM fungal species rather than between AM fungal species.

We find both ecological (i.e. changes in relative abundance of AM fungal species) and evolutionary (i.e changes in genetic composition within AM fungal species) dynamics operating concurrently within the plant – AM fungal symbiosis. We find strong evidence that individual AM fungal species, and genotypes within species, vary in their response to plant species, and that patterns of AM fungal divergence were structured by both plant phylogeny and plant traits governing life history variation. AM fungal composition generally improved host fitness, i.e. mycorrhizal feedbacks were generally positive. However, there was substantial variation in strength of mycorrhizal feedbacks, with variation in plant life history predicting stronger positive feedbacks, while variation due to plant phylogeny did not. Moreover, evolutionary changes within individual AM fungal species were the major driver of positive feedback rather than ecological changes in AM fungal species composition. Given the evidence that AM fungal diversity and composition can be critical to plant community and terrestrial ecosystem function [5–9], our work suggests that improved understanding of both evolutionary and ecological dynamics of AM fungi are necessary for optimal management of terrestrial ecosystems and projections of ecosystem response to anthropogenic disturbance.

## Materials and Methods

A common community of AM fungi made up of seven cultured isolates derived from tallgrass prairie (*Claroideoglomus lamellosum, Claroideoglomus claroideum, Entrophospora infrequens, Funneliformis mosseae, Racocetra fulgida, Cetraspora pellucida, Acaulospora spinosa*) was distributed into replicate mesocosms. The mesocosms were planted in the greenhouse with one of 38 grassland plant species from the prairie region chosen to provide a high degree of both phylogenetic diversity as well as diversity in life history traits associated with plant successional stage (S1 and S2 Figs). AM fungal composition in mesocosms was allowed to differentiate in response to host species over two growing seasons (2017 and 2018) and the composition of the AM fungal community was sampled after each growing season. In both years plants were grown in the greenhouse for the entire growing season (March to September) and were harvested in the fall. Plants were grown in two-gallon pots with four individuals of a species to a pot in the first year. After the first year, half of the soil in each pot was retained as inocula for the same host species and same replicate in year 2. Soil samples were taken after each growing season for determination of AM fungal composition using next-gen Illumina™ sequencing. We assumed over-representation of sequences as indicative of positive fitness impacts in plant treatments and under-representation indicative of negative fitness impacts. In year 3 we conducted a greenhouse feedback experiment where each soil community, trained by association with specific plant species, was used as inocula for other plants in the experiment. These were grown with a single host plant individual in 1-liter deep pots and inoculated with 100 ml of soil from the year 2 mesocosms. From the plant biomass data we were able to calculate feedback coefficients.

### Meta-analysis Methods

The seven AM fungi isolates used in this experiment were used previously in three separate studies testing their impact on early and late successional plant species: one reported in [45] and two in [28]. We conducted a meta-analysis of these studies to obtain estimates of the growth promotion for each isolate to plant species of different plant phylogenetic groups (grass, asters, and legumes) and from different life histories/origin groups (early successional native, non-native, and late successional native plants). We calculated the log mycorrhizal responsiveness (LRR) using equation (1) where <u>*x*</u>_*inoc*_ and <u>*x*</u>_*ctrl*_ are mean plant biomass for the inoculated treatments and sterile controls, respectively. The sampling variance (*σ*_2_) of LRR was estimated using equation (2) where 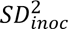 and 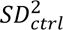 are the standard deviations and *n*_*inoc*_ and *n*_*ctrl*_ are the sample sizes of the treatment and controls respectively (Hedges et al., 1999; Hoeksema et al., 2018).

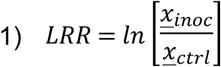

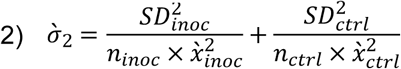

The analysis was conducted using the metafor package in R [61]. Moderators were the host plant life history, host plant phylogenetic group, the AM fungal isolate used, and the interaction of the isolate used and plant life history and phylogenic group. Random effects included individual experiments and plant host species. The model was estimated using the restricted maximum-likelihood (“REML”) method. Marginal means were estimated for the interaction terms using the emmeans package [62]. The metafor model was converted to a reference grid for compatibility with the emmeans package using the qdrg function.

We also tested whether plant life history predicted plant response to differences in AM fungal diversity of inocula. Only the two studies in Cheeke et al. (2019) included multi-species with single AM fungal species inoculations. Following the approach developed in Magnoli and Bever (2023), we calculated LRR in this analysis as the ratio of multiple species inocula mean plant biomass <u>*x*</u>_*multi*_ to the mean plant biomass averaged across all single inocula <u>*x*</u>_*mean*_. We also calculated the ratio of <u>*x*</u>_*multi*_ to the mean plant biomass of the single most beneficial AM fungal inocula <u>*x*</u>_*most*_. For each of these tests we constructed measures of variance by modifying equation 2.

### Bioinformatics Methods

To measure the changes in relative abundances of AM fungal species 0.2 g of soil was retained from each pot for Illumina™ sequencing. Extraction was performed on 311 samples using the Qiagen DNeasy Powersoil Kit. One sample of the 311 was lost during storage in both years. DNA was amplified using the LROR and FLR2 primers [63,64]. Volumes of 12.5μL of Phusion® Hot Start Flex DNA Polymerase master mix, 0.5μL of each primer, and 10.5μL of MilliporeSigma™ Direct-Q™ 3 type I water per sample were combined with 1μL of extracted DNA. PCR was conducted using one 5 minute cycle at 94.0 C, thirty-five cycles alternating between 94.0 C for 30 seconds, 48.0 C for 30 secs, 72.0C for 30 seconds and 72.0 C for 10 minutes.

Bioinformatics was conducted using the Dada2 and Qiime 2 pipelines as described in [40–42]. Primers were removed using Cutadapt. Using Dada2 forward reads were truncated to a length where 95% of base pair Phred quality scores (Q scores) had a greater than 99% base call accuracy, with forward reads truncated to a length of 190bp and reverse reads to a length of 140bp. Forward and reverse reads were merged using the “justconcatonate” flag in Dada2 and chimeras were removed. Taxonomy was assigned by building a phylogenetic tree of the amplicon sequence variants (ASVs) using RAxML together with an included library of known AM fungi species [40,41,43,65,66]. All AM fungi identified sequences were blasted against the reference library. Several ASVs that were placed inside AM fungi had unacceptably long branch lengths and were suspected of being non homologous gene regions that were erroneously being placed in the phylogeny [40]. Those ASVs were excluded from the analysis. All background soil was sterilized. Therefore, the ASVs that were not identified as one of the seven inoculated species were assumed likely to be legacy DNA from dead organisms in the background soil and were also not included in the statistical analysis. ASVs were assigned a species taxonomy if they clustered within known species sequences from the reference library.

### Plant Phylogenetic Methods

A phylogeny was constructed for all plants in the study to determine phylogenetic distance. Sequences were obtained for the rbcL gene for all 38 species of plants from the NCBI GenBank database. A constraint tree was obtained using the Phylomatic “Tree of Trees” software using the slik megatree [67,68]. Tree construction was conducted using RaXML [66].

### Statistical Methods

ASV abundances for inoculated AM fungal species were centered log transformed and a PerMANOVA analysis was conducted using the adonis function in the R package vegan [69]. The predictors used were host plant life history, host plant phylogenetic group, and the interaction of host plant life history with host plant phylogenetic group. Phylogenetic groups are characterized as lilies (Amaryllidaceae and Commelinaceae), grasses (Poaceae), legumes (Fabacea), milkweeds (Apocynaceae), mints (Lamiaceae and Plantaginaceae), and asters (Apiaceae and Asteraceae).

Using a categorical factor, such as phylogenetic group, to analyze the effect of phylogenetic relationships is a limited approach. We used a method to include phylogenetic distances as a random effect to preserve the information included in the full phylogenetic tree. When visualizing the phylogenetic relationships and AM fungal composition, there are hints at relationships that would be difficult to test using more traditional statistical methods (Fig 2). A phylogenetic generalized mixed model (PGLM) was fitted using the phyr R package [70,71]. Host plant successional status, and the interaction between them were modeled as fixed effects. Block and the log transformed sequences were included alongside phylogenetic distance and species as random effects. As a univariate analysis each AM fungal species’ ASVs were grouped, and their abundances logit transformed. We also assessed Shannon’s Diversity and the logit-transformed proportion of the inoculated AM fungi over the total number of ASVs. The reason for analyzing this proportion is that as the legacy background DNA would not be growing in response to the treatments, the proportion of inoculated AM fungi could be used as a proxy for changes in fungal density.

We purposely included full factorial tests of plant species and training plant species for subsets of the 38 plant species. Each subset was treated as a block, and we attempted to maximize comparisons across the phylogeny, while maintaining a relatively even number of both early and late successional plants (S12 Fig). Within these sub-set pairs of plant species, we estimated pairwise feedbacks that govern the influence of mycorrhizal fungal dynamics on plant-plant interactions. Pairwise feedbacks are measured by the interaction coefficient (*I*), as derived from the models of interguild frequency dependence and host-microbiome feedback. It is calculated according to equation (3) [19,47]. Where *F*_*Aα*_ is the fitness of plant *A* in soil *α*, *F*_*Aβ*_ the fitness of Plant *A* in soil *β*, *F*_*Bα*_ the fitness of plant *B* in soil *α*, and *F*_*Bβ*_ the fitness of plant *B* in soil *β*. In this experiment we used plant biomass as our measure of plant fitness. If this coefficient is positive then the plants are experiencing positive feedback, and if the coefficient is negative the plants are experiencing negative feedback. We used a type III ANOVA analysis to test the effect on the interaction coefficient from host plant and training plant life history and the interaction between them.

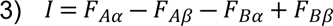

Dissimilarity between species compositions and genetic compositions within each AM fungal species was calculated using Aitchison distances (Euclidean distances based on center log transformed counts). For species composition all ASVs assigned to a single AM fungal species were first summed. For genetic change dissimilarity, separate dissimilarity matrices were calculated for all ASVs characterized as belonging to each AM fungal species. These dissimilarities were used in the PerMANOVA analyses. The list of pairwise dissimilarities derived from these matrices were used in regression analysis against feedback strength (interaction coefficients)

## Acknowledgments

We are grateful for the assistance of Sheena Parsons, Alice Tipton, Liz Koziol, Terra Lubin, Alexa Phillips, Reagan Smith, Kristen Mecke, Kelly Chesus, and the many undergrad technicians for assistance in project initiation, inocula preparation, and the planting, harvesting, and processing of the feedback experiment. We are also grateful for financial support from NSF grants DEB-1556664, OIA-1656006, DBI-2027458, BII-2120153, and DBI-2153040.

## Supporting information

**S1 Fig. Host Plant Characteristics.** Plant species were selected to represent a phylogenetically diverse group of early and late successional plants. Plants were selected from 5 large phylogenetic groups; grasses (Poaceae), legumes (Fabaceae), asters (Asteraceae and Apiaceae), mints (Lamiaceae and Plantaginaceae), milkweeds (Apocynaceae and Asclepiadaceae), and lilies (Liliaceae and Commelinaceae).

**S2 Table. Plant Species Used.**

**S3 Table. PerMANOVA Results for AM Fungi Species.** To test for overall differences in AM fungal composition with plant life history and plant family, we conducted a PerMANOVA on centered log ratio (CLR) transformed counts of ASVs grouped by AM fungal species. This analysis reveals statistically significant effects on AM fungal species composition of plant phylogenetic group in both years (p≤0.001, p≤0.001). We also tested for overall differences in AM fungal composition with plant life history and plant family using the PerMANOVA approach. This analysis reveals statistically significant effects on AM fungal species composition of plant life history on AM fungal species composition in year two (p<0.001)

**S4 Table. PerMANOVA Results for AMF ASVs Variation within Species.** As AM fungi are known to have substantial variability in rDNA composition within individual multinucleate cells, we also test for changes in ASV composition within individual AM fungal species using a PerMANOVA approach. For E. infrequens and Cl. claroideum we find evidence of change in genetic composition with plant phylogenetic group in both year one (p≤0.001, p≤0.01) and year two (p≤0.1, 0.01). Genetic composition of Cl. lamellosum and Ce. pellucida showed significant responses of plant phylogenetic group in year 1 (p≤0.001, p≤0.001), while genetic composition of F. mosseae and R. Fulgida had significant effects in year 2 (p≤0.01, p≤0.001). The genetic composition of F. mosseae, Cl. claroidium, and R. fulgida showed an effect of host plant life history on intraspecies ASV variation in year 2 (p<0.001, p<0.01, p<0.05).

**S5 Table. PGLMM Results for AM Fungi Species.** Phylogenetic generalized linear mixed model (PGLMM) results for relative proportions of each AM fungal species (combined ASV counts), as well as Shannon Diversity and Logit transformed density estimates.

**S6 Fig. Marginal Means of Relative Abundance by Plant Life History.** These are the estimated marginal means of AM fungal relative abundances when host plants were early or late successional. AM fungal species are arranged top to bottom in order of most to least beneficial. S7 Fig. Meta-Analysis Results. The marginal means for the log response ratio’s (LRT) of each of the seven AM fungal species used in this study. The LRT is the natural log of the ratio between how large plant species grew with AM fungal inoculation versus how they grew in a sterile control. These are derived from a meta-analysis of three previous experiments using these same cultures. (A) shows late successional plant benefits when grown in association with E. infrequens, Cl. lamellosum, F. mosseae, and Cl. claroideum. (B) shows the marginal means for the log response ratio’s (LRT) of each of the seven AM fungal species used in this study by host plant phylogenetic group. Asteraceae experienced significantly greater growth than Poaceae in association with all Claroideoglomus species (including E. infrequens). We also saw significant differential greater growth in Liliaceae over Poaceae as well as Liliaceae over Asteraceae in several AM fungal species.

**S8 Fig. LRR by Life History of AM Fungal Mixtures to Single Simbiont.** These are the log response ratios of the mean plant biomass for plants inoculated with a mixture of AM fungi to the mean plant biomass representative of the average single symbiont and to the most effective single symbiont respectively. Each panel breaks our results depending on if the host plant was an early or late successional.

S9 Fig. Histogram of mycorrhizal feedback measured as pairwise interaction coefficients. This shows counts of all paired comparisons with both the median and mean being significantly positive (p = 0.003, p≤ 0.001).

**S10 Fig: Feedbacks by Plant Phylogenetic Group.** Mycorrhizal feedback, as measured by the pairwise interaction coefficient, compared across phylogenetic group pairings. Feedback between individual pairs of plant species are indicated with dots, with pairwise feedback that was significantly different from zero indicated in black. We note that pairwise mycorrhizal feedbacks ranged from significantly negative to significantly positive. On average, we observed significantly positive mycorrhizal feedback between two legume species (p≤0.01) and significant negative mycorrhizal feedback between aster and grass species (p≤0.05).

**S11 Fig: Mycorrhizal Feedback Strength Predicted by Genetic Dissimilarity of AM fungi for All AM fungal Species**

Regressions of strength of pairwise feedback against measures of AM fungal genetic dissimilarity. Pairwise feedback was significantly predicted by genetic dissimilarity of two AM fungal species, *E. infrequens* (p<0.001) and *Cl. lamellosum* (p<0.001).

**S12 Fig. Feedback Experimental Design.** The experimental design for the feedback experiment included 5 subset groups of plants where all plant pairings were made in a fully factorial design. Each group also included early and late successional plants. When these pairings are arranged phylogenetically it becomes clearer that we also have a good representation of species pairs across the plant phylogeny. This allows us to test pairwise feedbacks, host plant characteristics, and plant phylogeny effects in a single experiment.

