## Supplemental figures and tables for "Plant phylogeny and life history predict AM fungal species and genetic composition, but only life history and genetic composition predict feedback": S1_Fig.docx

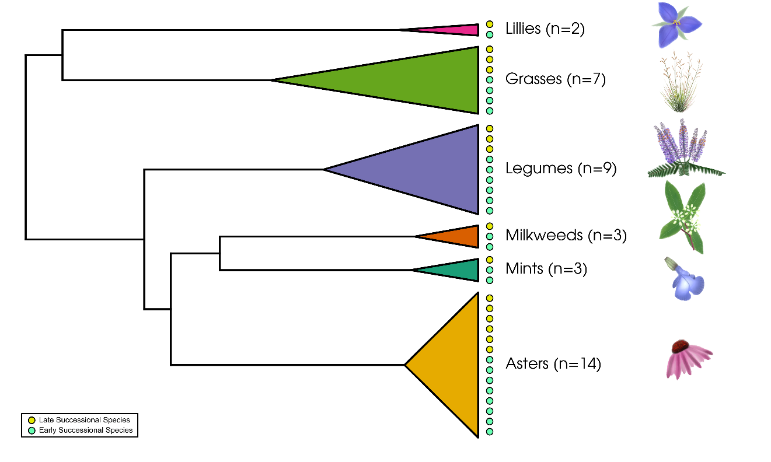

**Supplement S1: Host Plant Characteristics**
Plant species were selected to represent a phylogenetically diverse group of early and late successional plants. Plants were selected from 5 large phylogenetic groups; grasses (Poaceae), legumes (Fabaceae), asters (Asteraceae and Apiaceae), mints (Lamiaceae and Plantaginaceae), milkweeds (Apocynaceae and Asclepiadaceae), and lilies (Liliaceae and Commelinaceae).
