## Supplemental figures and tables for "Plant phylogeny and life history predict AM fungal species and genetic composition, but only life history and genetic composition predict feedback": S2_Table.docx

| **Supplement S2. Plant Species Used** | | | | | | |
| --- | --- | --- | --- | --- | --- | --- |
| Species | Family | Status |  | Species | Family | Status |
| Allium cernuum | Amaryllidaceae | late |  | Chamaecrista fasciculata | Fabaceae | early |
| Eryngium yuccifolium | Apiaceae | late |  | Desmanthus illinoensis | Fabaceae | early |
| Apocynum cannabinum | Apocynaceae | early |  | Desmodium illinoense | Fabaceae | early |
| Asclepias syriaca | Apocynaceae | early |  | Lespedeza capitata | Fabaceae | early |
| Asclepias tuberosa | Apocynaceae | late |  | Lespedeza cuneata | Fabaceae | early |
| Ambrosia artemisiifolia | Asteraceae | early |  | Melilotus officinalis | Fabaceae | early |
| Conyza canadensis | Asteraceae | early |  | Amorpha canescens | Fabaceae | late |
| Coreopsis tinctoria | Asteraceae | early |  | Baptisia alba | Fabaceae | late |
| Erechtites hieraciifolius | Asteraceae | early |  | Dalea purpurea | Fabaceae | late |
| Eupatorium altissimum | Asteraceae | early |  | Monarda fistulosa | Lamiaceae | early |
| Helianthus annuus | Asteraceae | early |  | Salvia azurea | Lamiaceae | early |
| Ratibida pinnata | Asteraceae | early |  | Pycnanthemum tenuifolium | Lamiaceae | late |
| Rudbeckia hirta | Asteraceae | early |  | Bromus inermis | Poaceae | early |
| Echinacea pallida | Asteraceae | late |  | Elymus canadensis | Poaceae | early |
| Liatris pycnostachya | Asteraceae | late |  | Panicum virgatum | Poaceae | early |
| Parthenium integrifolium | Asteraceae | late |  | Setaria faberi | Poaceae | early |
| Rudbeckia subtomentosa | Asteraceae | late |  | Andropogon gerardii | Poaceae | late |
| Silphium laciniatum | Asteraceae | late |  | Bouteloua curtipendula | Poaceae | late |
| Tradescantia ohiensis | Commelinaceae | early |  | Schizachyrium scoparium | Poaceae | late |
