## Supplemental figures and tables for "Plant phylogeny and life history predict AM fungal species and genetic composition, but only life history and genetic composition predict feedback": S3_Table.docx

| **Supplement S3: PerMANOVA Results for AM Fungi Species** | | | | | | | | |  |
| --- | --- | --- | --- | --- | --- | --- | --- | --- | --- |
|  | Year 1 | | | |  | Year 2 | | | |
|  | Df | R2 | Pr(>F) |  |  | Df | R2 | Pr(>F) |  |
| Plant life history | 1 | 0 | 1 |  |  | 1 | 0.03 | 0 | ٭٭٭ |
| Plant group | 5 | 0.1 | 0 | ٭٭٭ |  | 5 | 0.11 | 0 | ٭٭٭ |
| Plant species | 31 | 0.28 | 0 | ٭٭٭ |  | 30 | 0.29 | 0 | ٭٭٭ |
| Seq depth | 1 | 0 | 0.83 |  |  | 1 | 0.01 | 0.05 | ٭ |
| Block | 3 | 0.02 | 0.18 |  |  | 3 | 0.01 | 0.75 |  |
| Residual | 106 | 0.59 |  |  |  | 104 | 0.54 |  |  |
| Total | 147 | 1 |  |  |  | 144 | 1 |  |  |
| *** p ≤ 0.001; ** p ≤ 0.01; * p ≤ 0.05; · p ≤ 0.1 | | | | | | | | | |

To test for overall differences in AM fungal composition with plant life history and plant family, we conducted a PerMANOVA on centered log ratio (CLR) transformed counts of ASVs grouped by AM fungal species. This analysis reveals statistically significant effects on AM fungal species composition of plant phylogenetic group in both years (p≤0.001, p≤0.001). We also tested for overall differences in AM fungal composition with plant life history and plant family using the PerMANOVA approach. This analysis reveals statistically significant effects on AM fungal species composition of plant life history on AM fungal species composition in year two (p<0.001**)**
