## Supplemental figures and tables for "Plant phylogeny and life history predict AM fungal species and genetic composition, but only life history and genetic composition predict feedback": S4_Table.docx

| **Supplement S4: PerMANOVA Results for AMF ASVs Variation within Species** | | | | | | | |
| --- | --- | --- | --- | --- | --- | --- | --- |
|  |  | Plant life history | Plant group | Plant species | Seq depth | Block | Residual |
| *E. infrequens* | Year 1 | 0 | 0.13 *** | 0.13 *** | 0.01 · | 0.02 | 0.46 |
|  | Year 2 | 0 | 0.04 · | 0.46 *** | 0 | 0.05 ** | 0.44 |
| *Cl. lammellosum* | Year 1 | 0 | 0.18 *** | 0.27 ** | 0.01 | 0.06*** | 0.48 |
|  | Year 2 | 0 | 0.02 | 0.42 *** | 0 | 0.06 ** | 0.5 |
| *F. mosseae* | Year 1 | 0.01 | 0.04 | 0.18 | 0.01 | 0.02 | 0.75 |
|  | Year 2 | 0.04 *** | 0.06 ** | 0.42 *** | 0 | 0.01 | 0.46 |
| *Cl. claroidium* | Year 1 | 0 | 0.06 * | 0.21 | 0.01 | 0.02 | 0.7 |
|  | Year 2 | 0.03 ** | 0.07 * | 0.24 | 0 | 0.01 | 0.65 |
| *R. fulgida* | Year 1 | 0 | 0.05 | 0.26 * | 0 | 0.01 | 0.67 |
|  | Year 2 | 0.02 * | 0.13 *** | 0.23 * | 0.01 | 0.03 | 0.58 |
| *Ce. pellucida* | Year 1 | 0 | 0.08 *** | 0.21 | 0.01 | 0.05 *** | 0.64 |
|  | Year 2 | 0 | 0.05 | 0.18 | 0 | 0.01 | 0.75 |
| *A. spinosa* | Year 1 | 0 | 0.03 | 0.21 | 0 | 0.02 | 0.74 |
| *** p ≤ 0.001; ** p ≤ 0.01; * p ≤ 0.05; · p ≤ 0.1 | | | | | | | |

As AM fungi are known to have substantial variability in rDNA composition within individual multinucleate cells, we also test for changes in ASV composition within individual AM fungal species using a PerMANOVA approach. For *E. infrequens* and *Cl. claroideum* we find evidence of change in genetic composition with plant phylogenetic group in both year one (p≤0.001, p≤0.01) and year two (p≤0.1, 0.01). Genetic composition of *Cl. lamellosum* and *Ce. pellucida* showed significant responses of plant phylogenetic group in year 1 (p≤0.001, p≤0.001), while genetic composition of *F. mosseae* and *R. Fulgida* had significant effects in year 2 (p≤0.01, p≤0.001). The genetic composition of *F. mosseae*, *Cl. claroidium*, and *R. fulgida* showed an effect of host plant life history on intraspecies ASV variation in year 2 (p<0.001, p<0.01, p<0.05).
