## Supplemental figures and tables for "Plant phylogeny and life history predict AM fungal species and genetic composition, but only life history and genetic composition predict feedback": S5_Table.docx

| **Supplement S5: PGLMM Results for AM Fungi Species** | | | | | | | |
| --- | --- | --- | --- | --- | --- | --- | --- |
|  |  | Host Life History | Host Species | Host Phylogentic | Block | Log Seq Depth | Residual |
| *E.  infrequens* | Year1 | 0.08 (0.21) | 0.00 (0.00) *** | 1.02 (0.60) *** | 0.00 (0.00) | 0.00 (0.00) | 2.84 (1.69) |
|  | Year2 | 0.05 (0.39) | 0.80 (1.83) *** | 1.63 (2.61) * | 0.00 (0.00) | 11.47 (6.93) | 0.24 (0.49) |
| *Cl. lammellosum* | Year1 | -0.04 (0.25) | 0.66 (0.38) *** | 0.37 (0.28) | 0.37 (0.28) * | 0.00 (0.02) | 4.57 (2.14) |
|  | Year2 | 0.05 (0.34) | 1.42 (2.58) | 0.00 (0.01) | 0.28 (1.15) | 11.17 (7.23) | 0.21 (0.46) |
| *F. mosseae* | Year1 | -0.18 (0.27) | 0.00 (0.00) | 0.04 (0.07) | 0.00 (0.00) | 2.38 (0.56) | 7.54 (2.75) |
|  | Year2 | -0.48 (0.28) · | 2.46 (4.10) *** | 0.00 (0.00) | 0.00 (0.00) | 1.50 (3.20) | 0.15 (0.38) |
| *Cl. claroidium* | Year1 | 0.12 (0.29) | 0.00 (0.01) | 0.28 (2.99) | 0.00 (0.03) | 10.73 (18.44) · | 0.03 (0.18) |
|  | Year2 | -0.54 (0.24) * | 0.58 (1.40) | 0.02 (0.28) | 0.04 (0.39) | 5.27 (4.23) | 0.29 (0.54) |
| *R.  fulgida* | Year1 | -0.30 (0.45) | 4.43 (3.40) *** | 0.21 (0.74) | 0.40 (1.02) | 9.93 (5.09) | 0.38 (0.62) |
|  | Year2 | -0.68 (0.36) · | 0.60 (1.73) *** | 0.87 (2.09) *** | 0.00 (0.00) | 11.78 (7.67) | 0.20 (0.45) |
| *Ce. pellucida* | Year1 | -0.17 (0.30) | 0.03 (0.06) *** | 0.79 (0.30) * | 0.80 (0.30) * | 0.00 (0.00) | 8.91 (2.98) |
|  | Year2 | -0.18 (0.14) | 0.35 (0.00) ** | 0.00 (0.00) | 0.00 (0.00) | 0.99 (1.69) | 0.35 (0.59) |
| *A. spinosa* | Year1 | -0.14 (0.24) | 0.00 (0.01) | 0.00 (0.76) | 0.02 (804.61) *** | 8.29 (14,907.21) *** | 0.00 (0.00) |
| Shannon Diversity | Year1 | -0.02 (0.03) | 0.00 (0.00) *** | 0.01 (0.47) ** | 0.00 (0.14) | 0.05 (0.89) | 0.06 (0.25) |
|  | Year2 | -0.09 (0.04) * | 0.04 (0.83) *** | 0.00 (0.00) | 0.00 (0.3) · | 0.02 (0.58) | 0.05 (0.23) |
| Logit  Density | Year1 | -0.16 (0.11) | 0.00 (0.01) *** | 0.34 (3.05) *** | 0.01 (0.6) | 0.68 (4.31) | 0.04 (0.19) |
|  | Year2 | -0.02 (0.21) | 1.28 (1.57) *** | 0.00 (0.00) | 0.19 (0.6) ** | 1.04 (1.41) | 0.52 (0.72) |
| *** p<0.001; ** p<0.01; * p<0.05; · p<0.1 | | | | | | | |

Phylogenetic generalized linear mixed model (PGLMM) results for relative proportions of each AM fungal species (combined ASV counts), as well as Shannon Diversity and Logit transformed density estimates.
