## Supplemental figures and tables for "Plant phylogeny and life history predict AM fungal species and genetic composition, but only life history and genetic composition predict feedback": S6_Fig.docx

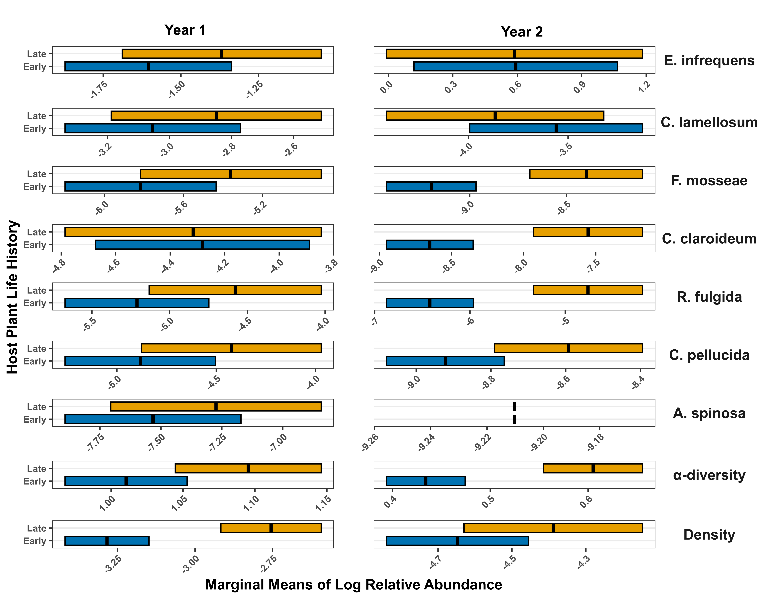

**Supplement S6: Marginal Means of Relative Abundance by Plant Life History**
These are the estimated marginal means of AM fungal relative abundances when host plants were early or late successional. AM fungal species are arranged top to bottom in order of most to least beneficial.
