## Supplemental figures and tables for "Plant phylogeny and life history predict AM fungal species and genetic composition, but only life history and genetic composition predict feedback": S7_Fig.docx

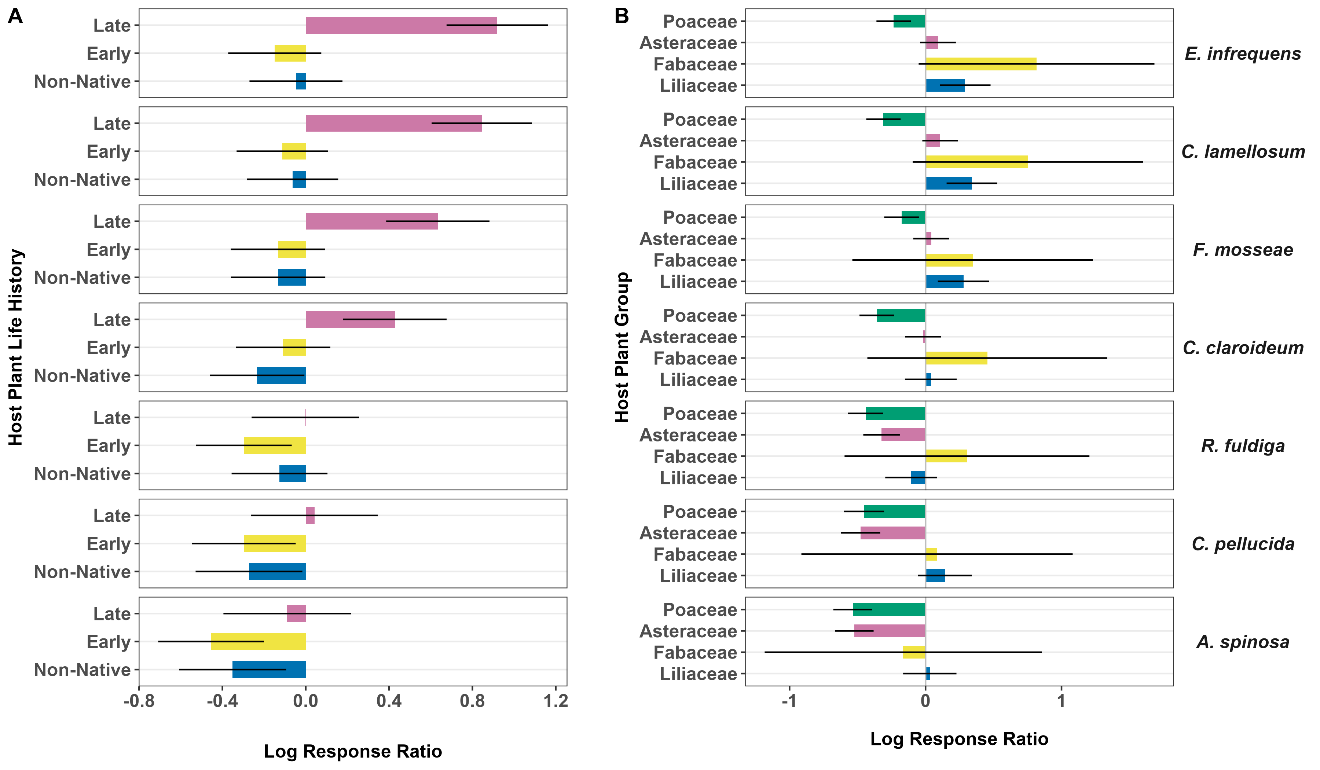

**Supplement S7: Meta-Analysis Results**
The marginal means for the log response ratio’s (LRT) of each of the seven AM fungal species used in this study. The LRT is the natural log of the ratio between how large plant species grew with AM fungal inoculation versus how they grew in a sterile control. These are derived from a meta-analysis of three previous experiments using these same cultures. (A) shows late successional plant benefits when grown in association with *E. infrequens*, *Cl. lamellosum*, *F. mosseae*, and *Cl. claroideum*. (B) shows the marginal means for the log response ratio’s (LRT) of each of the seven AM fungal species used in this study by host plant phylogenetic group. Asteraceae experienced significantly greater growth than Poaceae in association with all Claroideoglomus species (including *E. infrequens*). We also saw significant differential greater growth in Liliaceae over Poaceae as well as Liliaceae over Asteraceae in several AM fungal species.
