## Supplemental figures and tables for "Plant phylogeny and life history predict AM fungal species and genetic composition, but only life history and genetic composition predict feedback": S8_Fig.docx

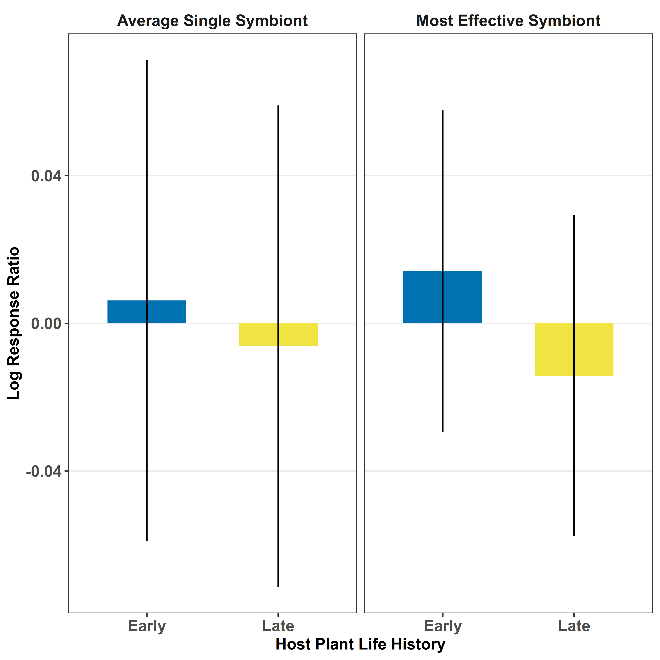

**Supplement S8: LRR by Life History of AM Fungal Mixtures to Single Simbiont**
These are the log response ratios of the mean plant biomass for plants inoculated with a mixture of AM fungi to the mean plant biomass representative of the average single symbiont and to the most effective single symbiont respectively. Each panel breaks our results depending on if the host plant was an early or late successional.
