## Supplemental figures and tables for "Plant phylogeny and life history predict AM fungal species and genetic composition, but only life history and genetic composition predict feedback": S9_Fig.docx

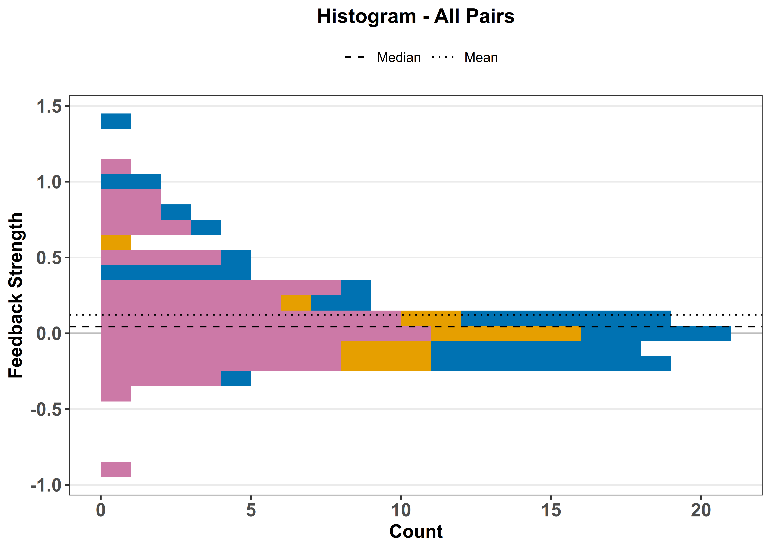

**Supplement S9:** Histogram of mycorrhizal feedback, measured as pairwise interaction coefficients. This shows counts of all paired comparisons with both the median and mean being significantly positive (p = 0.003, p≤ 0.001).
