## Supplemental figures and tables for "Plant phylogeny and life history predict AM fungal species and genetic composition, but only life history and genetic composition predict feedback": S10_Fig.docx

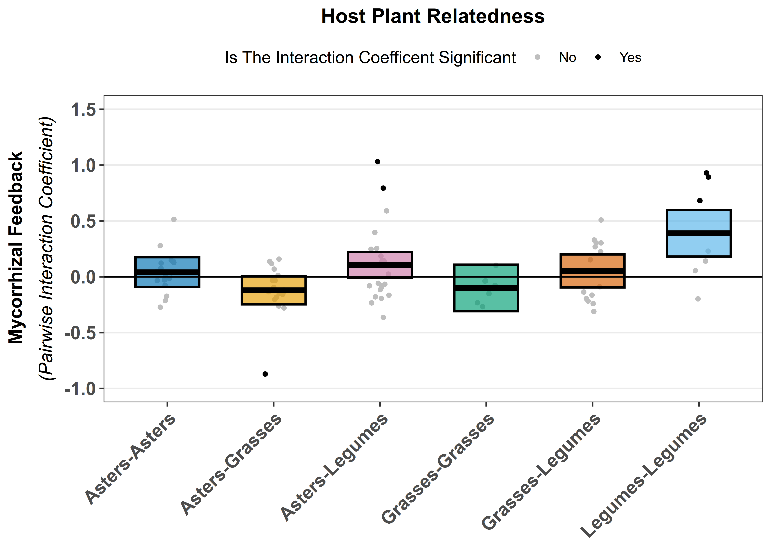

**Supplement S10: Feedbacks by Plant Phylogenetic Group**

Mycorrhizal feedback, as measured by the pairwise interaction coefficient, compared across phylogenetic group pairings. Feedback between individual pairs of plant species are indicated with dots, with pairwise feedback that was significantly different from zero indicated in black. We note that pairwise mycorrhizal feedbacks ranged from significantly negative to significantly positive. On average, we observed significantly positive mycorrhizal feedback between two legume species (p≤0.01) and significant negative mycorrhizal feedback between aster and grass species (p≤0.05).
