## Supplemental figures and tables for "Plant phylogeny and life history predict AM fungal species and genetic composition, but only life history and genetic composition predict feedback": S11_Fig.docx

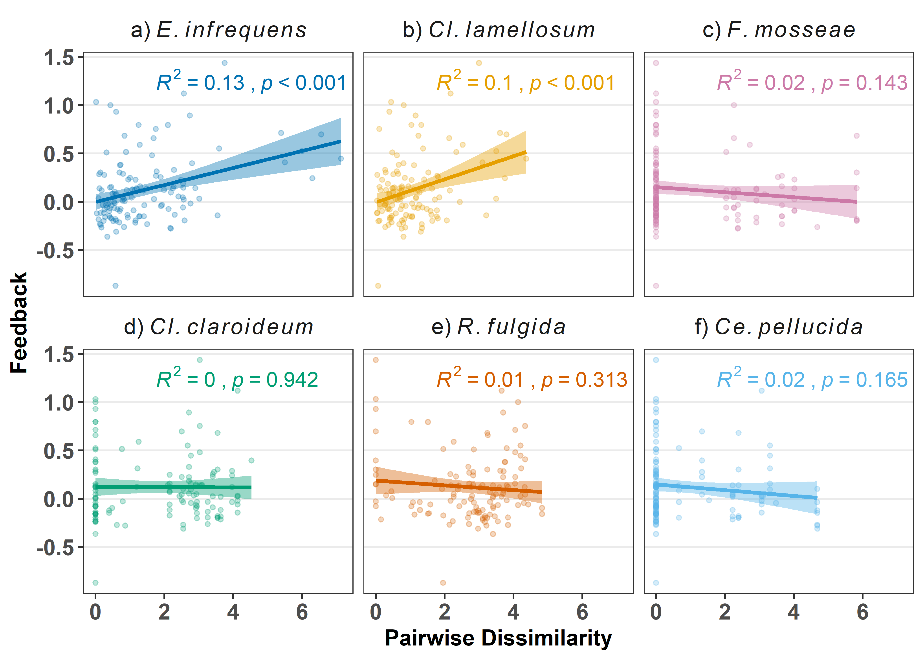

**S11 Fig: Mycorrhizal Feedback Strength Predicted by Genetic Dissimilarity of AM fungi for All AM fungal Species**

Regressions of strength of pairwise feedback against measures of AM fungal genetic dissimilarity. Pairwise feedback was significantly predicted by genetic dissimilarity of two AM fungal species, *E. infrequens* (p<0.001, F=17.78=, R^2^=0.13) and *Cl. lamellosum* (p<0.001, F=13.27, R^2^=0.1). Other species were not significant, including *F. mosseae* (p=0.143, F=2.175, R^2^=0.02), *C. claroideum* (p=0.942, F=0.005, R^2^=0), *R. fulgida* (p=0.313, F=1.026, R^2^=0.01), and *Ce. pellucida* (p=0.165, F=1.952, R^2^=0.02)
