## Supplemental figures and tables for "Plant phylogeny and life history predict AM fungal species and genetic composition, but only life history and genetic composition predict feedback": S12_Fig.docx

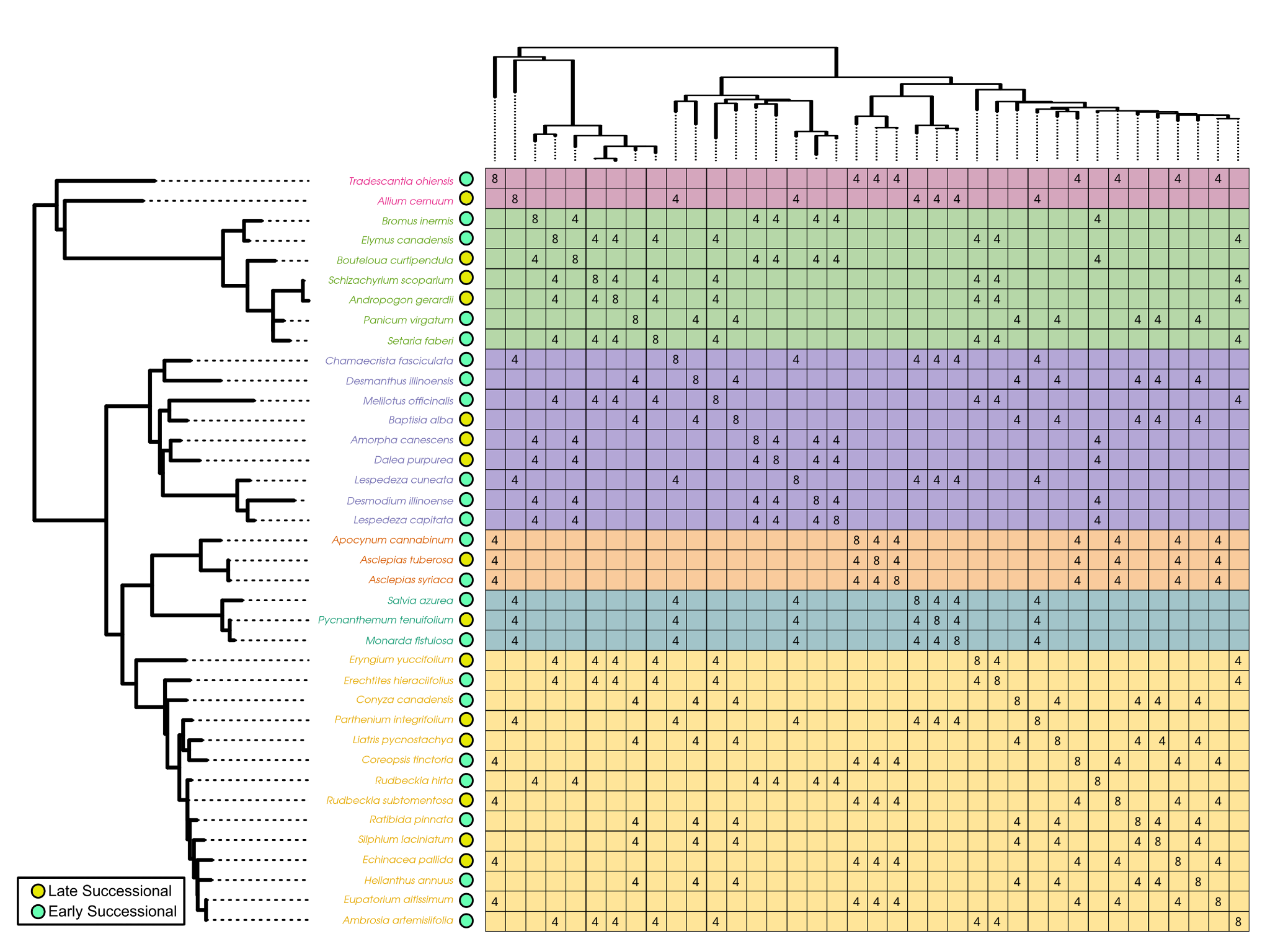

**Supplement S12: Feedback Experimental Design**

The experimental design for the feedback experiment included 5 subset groups of plants where all plant pairings were made in a fully factorial design. Each group also included early and late successional plants. When these pairings are arranged phylogenetically it becomes clearer that we also have a good representation of species pairs across the plant phylogeny. This allows us to test pairwise feedbacks, host plant characteristics, and plant phylogeny effects in a single experiment.
